# Host control of *Mycobacterium tuberculosis* infection is not influenced by the gut microbiome

**DOI:** 10.64898/2026.08.05.743129

**Authors:** Carolina Agudelo, Amir Balakhmet, Samuel B. Berry, Douglas M. Fox, Sarah A. Stanley, Ashley R. Wolf

## Abstract

Tuberculosis (TB) is a life-threatening disease with heterogenous presentation. Approximately one-quarter of the global population is infected with *Mycobacterium tuberculosis* (Mtb), yet a much smaller fraction develops active TB disease. Host genetics, immune system function, and environmental factors have all been implicated in susceptibility to Mtb, yet no one factor fully explains TB heterogeneity. Strikingly, many of these same factors are linked to gut microbiome composition, which is intimately linked to systemic development of the immune system. Antibiotic treated mouse models suggest that increased gut microbiota diversity is protective against Mtb infection. In contrast, *Helicobacter hepaticus* colonization is correlated with exacerbated Mtb burden. However, antibiotics can have both microbial and nonmicrobial targets and studies to date have not deconvoluted these effects. Focused testing of specific microbiome members has been impossible without a gnotobiotic model for Mtb. Here, we develop the first gnotobiotic mouse model for Mtb infection and test how microbial diversity in the gut microbiome impacts host susceptibility to Mtb. Surprisingly, after intranasal challenge with Mtb, germ-free wild type mice had no difference in lung burden when compared to mice born with either a defined gut microbiome community (OMM-12) or a diverse, conventional microbiota. *H. hepaticus* gut colonization of OMM-12 and conventional mice also did not impact Mtb burden in the lungs in this controlled setting. *H. hepaticus* colonization of the gut did perturb lung immune responses associated with TB infection control. CD4+ T cells were decreased, CD8+ T cells were increased, and IL-6 production was decreased. While the gut microbiome may yet play a role in immune compromised mouse models or human disease, comparing drastically different gut microbiotas in gnotobiotically controlled C57BL/6 mice did not yield any evidence of alteration in Mtb lung burden. The host’s unique immune response to Mtb may in part make the pathogen resistant to immune disruption caused by gut microbiome changes.

## Introduction

Cross-talk between the gut and the lung is proposed to modulate disease severity in a range of infectious and noninfectious diseases, ranging from COVID-19, to cancer. Disruption of the gut microbiome is hypothesized to increase susceptibility to the lung pathogen *Mycobacterium tuberculosis* (Mtb)^1^. Mtb infects approximately 10 million people annually and causes the deadly lung disease tuberculosis (TB)^2^. While human TB presentation varies from asymptomatic to fatal, no one factor can fully explain the heterogeneity observed^3^. Changes to the gut microbiome are known to influence metabolism and immune responses at distal sites including the lung, leading us to hypothesize that microbiome variation may contribute to TB disease susceptibility in mice and humans.

Human microbiome studies in tuberculosis patients are limited but suggest that pulmonary tuberculosis is associated with gut microbiome dysbiosis. TB patients were found to have lower overall diversity, characterized by an increase in abundance of opportunists like *Enterococcus*, *Rothia*, and *Clostridia* and a decrease in short-chain fatty acid (SCFA) producers like Bifidobacterium^4–6^.

Disruption of the gut microbiome has been linked to susceptibility to lung pathogens including *Acinetobacter baumanii*, *Streptococcus pneumoniae*, and *Klebsiella pneumoniae*^7–9^. The protection offered by the gut microbiome to the host can occur directly, through competition in the gut, and indirectly, through modulation of the immune system^10^. For instance, commensal colonization in the intestine has been shown to promote resistance to *S. pneumoniae*, *K. pneumoniae* and *S. aureus* lung infections indirectly through induction of type 3 immune responses^8,11^.

Animal models exploring the role of the gut microbiome in TB have relied on antibiotic-treated mouse models. The use of broad-spectrum antibiotics in mice results in increased susceptibility to Mtb, enhancing inflammatory responses in the lungs and exacerbating pathology and Mtb lung burden^1,12,13^. Additionally, antibiotic-induced intestinal dysbiosis has been associated with impaired function of alveolar macrophages in the context of Mtb immunity^14^. However, antibiotic perturbations of the gut microbiome can be difficult to interpret because antibiotics may have direct and indirect impacts on the host. Colonization with *Helicobacter hepaticus* was also shown to impact immune response to Mtb in an immunocompromised model, leading to a slightly higher burden of disease in the lungs^15^.

The role of the gut microbiome in susceptibility to Mtb remains underexplored and challenging to interpret using current antibiotic-treated mouse models. We sought to understand the role of the gut microbiome in the context of Mtb infection using germ-free and gnotobiotic mouse models to enable precise manipulation of the gut microbiome. Here we report techniques for the use of gnotobiotic mice to maintain gut sterility for the duration of Mtb infection in a Biosafety Level 3 (BSL3) facility. We see that infection causes changes to the microbiome of conventional mice, consistent with observations in human TB studies^4–6^. Surprisingly, diversity of the gut microbiome does not influence host control of Mtb in this model. We further found that introduction of the pathobiont *Helicobacter hepaticus* remodels the lung immune environment but does not impact Mtb lung burden. Our findings speak to the importance of controlled gnotobiotic models and suggest that Mtb expansion in the lung is not impacted by gut-induced immune changes.

## Methods

### Bacterial cultures and strains

All OMM-12 strains used in this study are listed in Supplementary Table 1. All OMM-12 strains were grown in accordance with DSMZ’s culturing specifications. *Helicobacter hepaticus* ATCC 51449 was grown on Columbia blood agar (10% defibrinated horse blood) supplemented with 1% BCD, vancomycin, and amphotericin B in a BD BBL™ GasPak™ jar with BD GasPak™ EZ Campy Container System Sachets (REF 260680) at 37 °C. The *Mycobacterium tuberculosis* strain H37Rv was used for all experiments. H37Rv was grown in Middlebrook 7H9 broth supplemented with 10% OADC and 0.05% Tween 80 at 37 °C. CFU was determined by plating bacteria in 7H10 solid media.

### Mouse husbandry

C57BL/6NTac (B6-GF) mice were obtained from Taconic Biosciences (Germantown, NY) and bred in-house in sterile isolators (CBC Model: 50302424122). Sterility was tested by culturing fecal pellets in LYBHI aerobically and anaerobically for 2 days. Each strain of the OMM-12 community (Supplemental Table 1) was grown according to the specifications provided by DSMZ. Each strain was pooled at equal OD600 for oral gavage of mice. C57BL/6NTac germ-free mice (Taconic Biosciences) were colonized with 200 ul of the OMM-12 cocktail once via oral gavage. One week post gavage, fecal samples were collected to confirm colonization using 16S amplicon sequencing. Once colonization was confirmed, the OMM-12 colonized mice were bred. Progeny of the colonized mice were kept on an Innovive Innorack IVC Mouse 3.5 using Innovive’s caging (M-BTM bottom, MVX3 containment lid). C57BL/6NTac murine pathogen free (MPF®) mice were obtained from Taconic Biosciences for each experiment. For all experiments, mice were co-housed in groups of 5 mice maximum per cage in Innovive ventilated caging and fed an autoclaved Teklad Global 18% Protein Extruded Rodent Diet - Sterilizable (Innotiv 2018SX).

All animal work was done in strict accordance with the recommendations of the Guide for the Care and Use of Laboratory Animals of the National Institutes of Health and the University of California, Berkeley regulations. Protocols were reviewed and approved by the Animal Care and Use Committee at the University of California, Berkeley (AUP 2021-01-13960).

### H. hepaticus colonization

*H. hepaticus* colonies were resuspended in Brucella broth supplemented with 10% FBS. Each mouse was orally gavaged with 100 µl of > OD600=1 at least 1 week prior to Mtb infection. OMM-12 mice were gavaged once, while MPF mice were gavaged twice every 2 days. To confirm colonization, a PCR was performed on fecal gDNA using Neubert et al.’s *H. hepaticus* primers^16^. Cycling conditions were as follows: 95 °C for 5 min, followed by 35 cycles of 95 °C for 30 s, 55 °C for 30 s, and 72 °C for 1 min. Final extension for 5 min at 72 °C.

### Mouse infections

Eight- to twelve-week old C57BL/6NTac mice (Taconic BioSciences) were infected intranasally with 50-100 CFU of *Mycobacterium tuberculosis* strain H37Rv in 40 uL of sterile PBS-Tween (0.05%). Initial doses were confirmed by whole lung CFU 1 day post infection. Infections proceeded for 28 days. Mice were then euthanized and lungs were harvested, reserving the smallest lobe for histology and the remaining lungs used for CFU enumeration and flow cytometry. Lungs were plated on 7H10 plates for 3 weeks. Data shown represent at least 3 independent experiments.

### Flow cytometry

Whole lungs were harvested into complete RPMI-1640 media supplemented with 10% FBS, 1 mM sodium pyruvate (Sigma Aldrich S8636), 0.01 mM HEPES, 1% Glutamax (Thermo Scientific 35050061), 1% non-essential amino acids (Gibco 11140–050), and 55 μM beta-mercaptoethanol (Gibco 21985–023) containing Liberase TM (Roche 5401127001) and DNase I (Roche 11284932001). Lungs were processed in gentleMACS™ C tubes (Miltenyi 130-096-334) using the gentleMACS™ Octo Dissociator, then strained through a 70 μm cell strainer to generate a single cell suspension. 1/50th of the total suspension was aliquoted for enumeration of bacterial burden.

For flow cytometry immunophenotyping and intracellular cytokine staining, 1/15th of the lung homogenate was restimulated *ex-vivo* with complete RPMI containing optimized concentrations of *Mycobacterium tuberculosis* Ag85b peptide pool (peptides&elephants LB02088), *Mycobacterium tuberculosis* ESAT6 peptide pool (peptides&elephants LB02113), vehicle control or eBioscience Cell Stimulation Cocktail (Invitrogen 00-4970-03) in the presence of eBioscience Protein Transport Inhibitor Cocktail (Invitrogen 00-4980-03) for 5 hours at 37°C at 5% CO2. Cells were then blocked with BD Mouse Fc Block (BD Bioscience 553142) for 20 minutes at room temperature, then stained with optimized concentrations of anti-CD3e BUV661 (BD Bioscience 750638), anti-Ly6C BUV563 (BD Bioscience 755198), anti-CD11c BUV496 (BD Bioscience 750450), anti-CD64 BV786 (BD Bioscience 569507), anti-CD11b BV605 (BioLegend 101257), anti-SiglecF PerCP-Fire806 (BioLegend 155536), anti-CD8a RB613 (BD Bioscience 571099), anti-F4/80 StarBright Blue 580 (BioRad MCA497SBB580), anti-MerTK FITC (BioLegend 151504), anti-NK1.1 PE-Cy7 (BioLegend 108714), anti-CD4 PE-CF594 (BD 562285), and anti-Ly6G APC-Cy7 (BioLegend 127623) for 20 minutes at room temperature. Cells were stained for viability using Aqua LIVE/DEAD (Invitrogen L34957) for 20 minutes at room temperature, then fixed and permeabilized using BD Cytofix/Cytoperm (554714) according to manufacturer’s instructions. Permeabilized cells were stained for intracellular markers and secreted cytokines using optimized concentrations of anti-Arginase-1 BUV805 (Invitrogen 368-3697-82), anti-IL-10 Pacific Blue (BioLegend 505020), anti-IL-6 BV650 (Invitrogen 41-670-6182), anti-iNOS PerCP-eFluor 710 (Invitrogen 46-5920-82), and anti-TNFa PerCP-Cy5.5 (BioLegend 506322) overnight at 4 °C. Pooled leftover homogenates were used for fluorescence-minus-one (FMO) controls.

Data was acquired on a Cytek Aurora spectral cytometer with 5-laser configuration and analyzed in SpectroFlo after spectral unmixing with autofluorescence extraction. UltraComp eBeads (Invitrogen 01-2222-42) individually stained according to manufacturer’s instructions with all antibodies in the panel were used as unmixing controls. Gating for markers was determined in reference to unstimulated/PMAi-stimulated wells as well as fluorescence-minus-one (FMO) controls.

### DNA isolation and 16S rRNA gene amplicon library preparation

One fecal pellet was added to a sterile 2 ml screw-top tube with 500 μl 0.1 mm zirconia/silica beads, 500 μl of Buffer A (200 mM Tris-HCl, pH 8.0/200 mM NaCl/20 mMEDTA), 210 μl of 20% SDS, and 500 μl of phenol/chloroform/IAA (pH 7.9, 25:24:1). The sample was then bead beaten on a Bullet Blender lite at the highest speed for 4 minutes. The sample was centrifuged at 13,000 rpm and 4 °C for 3 minutes. Samples were then boiled for 1 hour at 80 °C. The aqueous phase was transferred out of the BSL3 to work in BSL1 conditions. 200 μl of the aqueous layer was then passed through Monarch’s Genomic DNA Purification Kit (Cat. No./ID: T3010S/L) per manufacturer’s protocol. gDNA was quantified using a Nanodrop and diluted to 10 ng/μl. 16S rRNA amplification was conducted using the KAPA HiFi Hotstart kit (Cat. No. 50-196-5215) and the primers targeting the V4 region as described by Gohl et al.^17^ (Supplemental Table 2). Cycling conditions were as follows: 95 °C for 5 min, followed by 25 cycles of 98 °C for 20 s, 55 °C for 15 s, and 72 °C for 1 min. Final extension for 10 min at 72 °C. A second PCR followed to add sample-specific indices and Illumina compatible flow cell adaptors (Supplemental Table 2). Cycling time was as follows: 95 °C for 5 min; ten cycles of 98 °C for 20 s, 55 °C for 15 s, and 72 °C for 1 min; and a final extension at 72 °C for 10 min. A left-side selection (0.8x) using SPRI beads followed the PCR using the manufacturer’s protocol. The PCR product was quantified using a Qubit and pooled for sequencing on an Illumina MiSeq to generate 2x250 bp paired end reads.

### Microbiome data analysis

The Quantitative Insights into Microbial Ecology 2 (Qiime2) pipeline version 2023.5.1^18^ was used for downstream analyses. Potential chimeric reads were removed using consensus-based methods. Amplicon sequence variants (ASVs) were inferred using DADA2^19^ and a phylogenetic tree was built using MAFFT^20,21^ alignment. Taxonomic assignment was performed using the SILVA138^22^ database. For alpha and beta diversity analyses, samples were rarefied to 2500 reads per sample using the Qiime2 pipeline.

### Statistical analysis

The metadata file and Qiime2 files were imported into R version 4.2.1 and merged into a single Phyloseq object. Analysis of composition of microbiomes with bias correction (ANCOMBC)^23^ was performed in R using the “ANCOMBC”^23,24^ package. Bacterial taxa were determined to be differentially present based on an adjusted p-value of < 0.05. Alpha diversity was calculated using Faith’s Phylogenetic Diversity (PD) and the Shannon Index, with between group significance determined using a Wilcoxon rank sum test. Using the Qiime2 package, beta diversity was calculated using the Weighted UniFrac distance matrix. PERMANOVA tests were done to see if the distributions were day post infection was used as the independent variable. PERMDISP was used to calculate whether the variances between two groups were significantly different. Plots were generated using the “ggplot2”^25^ package and base R functions and edited in Adobe Illustrator. Specific statistical tests and sample sizes can be found in the figure legends. Data were analyzed using GraphPad Prism 10. Flow cytometry data was analyzed using SpectroFlo V3.3.0 (Cytek).

### Data availability

Sequencing reads will be available through the NCBI Short Read Archive (SRA) under BioProject PRJNA1499356.

## Results

### Establishing a gnotobiotic model for tuberculosis

To test the role of the gut microbiome in Mtb susceptibility, we established a gnotobiotic model for Mtb infection. To our knowledge, no one has tested the host response to Mtb in either a germ-free or gnotobiotic mouse model. Germ-free and gnotobiotic mice are housed in gnotobiotic isolators to maintain sterility. Establishing and maintaining a gnotobiotic isolator is challenging in a biosafety level 3 (BSL3) facility, which is required for Mtb experiments. Therefore, we first sought to optimize BSL3 compatible germ-free and gnotobiotic mouse husbandry.

We used an Innovive Innorack IVC Mouse 3.5 Hepa-filtered ventilated housing system for BSL3 containment and maintenance of sterility. This system has been used by other research groups to study gnotobiotic mice in non-BSL3 conditions^26^. Mtb infected cages were maintained under negative pressure to prevent the escape of aerosolized Mtb consistent with BSL3 safety procedures. In contrast, standard gnotobiotic work is typically done under positive pressure. To ensure that the negative pressure cages remained a closed and sterile environment, we sealed all gaps between the lid and water bottle interface using autoclaved vinyl tape. Autoclaved water and diet were also provided to these mice. Cage changes within the biosafety cabinet and daily health checks were performed by a single investigator to ensure sterile protocol when cages were opened within an animal handling biological safety cabinet. With these adjustments and standard BSL3 procedures, germ-free status was maintained for four weeks. Sterility was confirmed for all experiments by collecting fecal, diet, and bedding samples for culturing in rich media in aerobic and anaerobic conditions^27^.

Aerosolization of Mtb is typically used in mouse models to mimic human transmission, but this route of administration presented a challenge when infecting germ-free mice^28^. Aerosolization uses a nebulizer and full-body inhalation exposure system (Glas-Col, Terre Haute, IN) with an aerosolization chamber that cannot be kept completely sterile. Instead, we performed intranasal infection of Mtb within a biosafety cabinet, which has been shown to result in comparable infection outcomes^29–31^. Tandem experiments using both methods of infection were carried out using germ-free mice and sterility was tested throughout the experiment. Fecal samples were taken at 0, 14, and 28 dpi for 16s rRNA amplicon sequencing to track microbiome changes in mice with communities and culture in anaerobic sachets to track germ-free sterility. As expected, aerosol infected mice exhibited microbial outgrowth from fecal cultures at 14 days post infection (dpi), whereas fecal samples of intranasally infected mice showed no microbial growth. Using intranasal administration of Mtb and the aforementioned negative pressure microisolator caging, gut sterility of germ-free mice was maintained for 28 days in the BSL3.

### Germ-free and conventional mice develop similar Mtb lung burden

To evaluate the role gut microbial diversity plays in host susceptibility to Mtb, we intranasally infected three groups of mice with different microbiomes with the Mtb H37Rv strain (target dose of 100 CFU/lung). We compared germ-free mice (GF), mice with the defined community OMM-12 (OMM), and conventional (Taconic murine pathogen-free, MPF) mice (Figure 1A). At 28 dpi, no differences in bacterial load as measured by colony forming units (CFU) were observed when comparing the lungs of all three communities across four independent experiments (Figure 1B, Supplementary Figure 1). We observed significant variability in bacterial burdens in the lung within an individual experiment consistent with what has been published for intranasal Mtb infections^30,31^. Histological analysis showed similar levels of immune infiltration surrounding lesions in infected mice from all three communities, consistent with equivalent bacterial loads and comparable immune responses (Figure 1C). Thus, diversity of the gut microbiome does not impact host susceptibility and immune response to Mtb in the lungs of infected mice.

**Figure 1.**
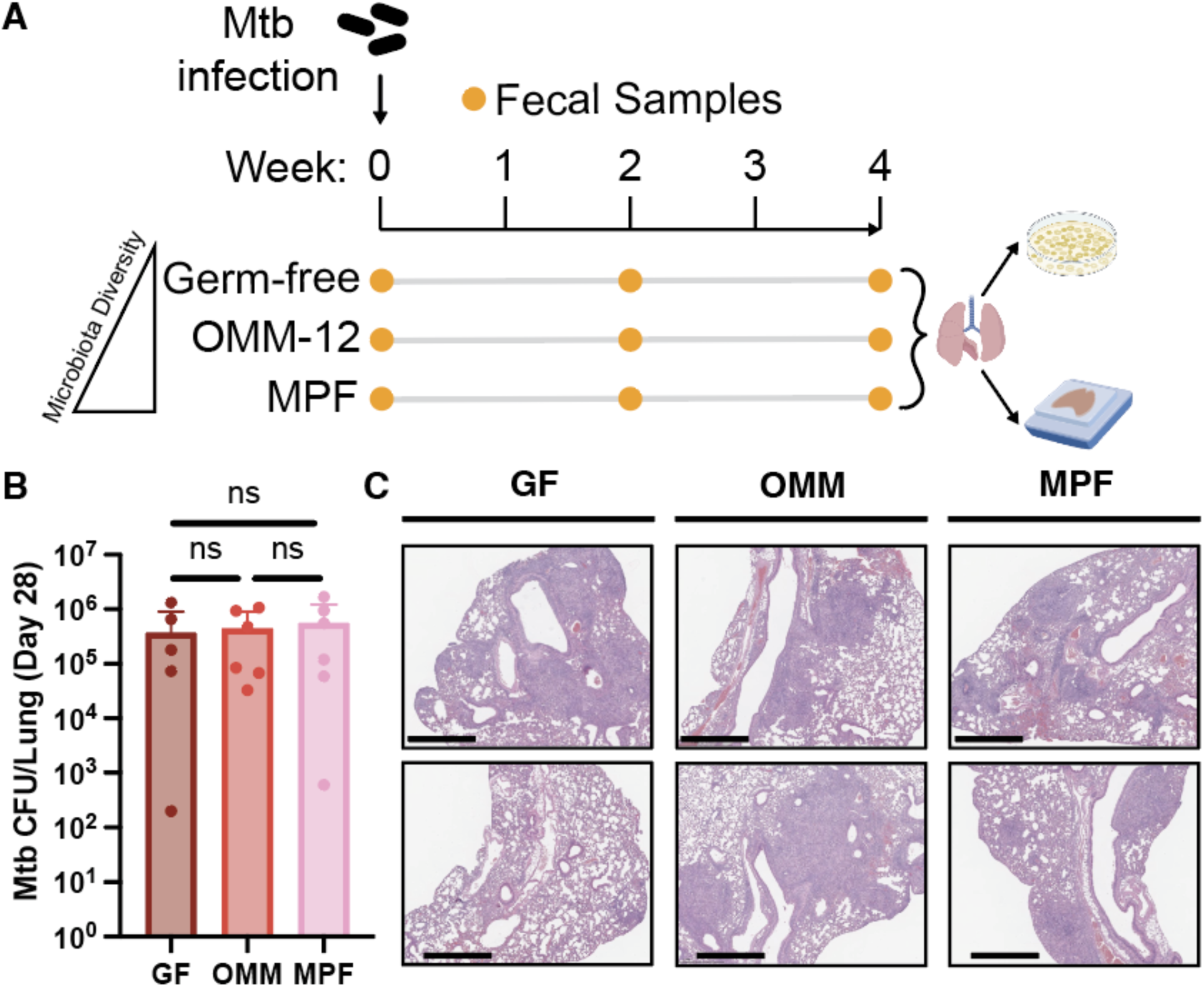
Gut microbiome diversity does not impact susceptibility to Mtb. (A) Experimental design of intranasal Mtb infections of germ-free (GF), OMM-12 (OMM), and conventional (MPF) mice. Fecal samples were collected at 0, 14, and 28 days post infection (DPI). Lungs were harvested at 28 dpi and used for plating for colony forming units (CFU) and histology. (B) Mtb burden in whole lung determined by CFU counts 3 weeks post harvest (NS, Mann-Whitney U test; bar plot representative of one replicate). (C) The smallest lobe was taken at harvest for histology and stained for H&E (bar = 1 mm).

### Mtb infection impacts gut microbiome composition

To determine whether Mtb infection causes alterations to the gut microbiome, we evaluated the composition of the gut microbiome of mice colonized with the defined OMM-12 community and their conventional counterparts (MPF) using 16S rRNA gene amplicon sequencing of genomic DNA (gDNA) isolated from fecal pellets. We found no evidence of a difference in alpha diversity when comparing OMM pre- and post-infection samples (Faith’s PD and Shannon’s Index, Supplementary Figure 2A,B). Analysis of beta diversity by Weighted UniFrac revealed that OMM pre- and post-infection samples are distinct from one another (Supplementary Figure 2C, p < 0.01, PERMANOVA; p = 0.413, PERMDISP). However, we observed stability in the relative abundance of individual OMM-12 members (Figure 2A), and identified no significantly changing taxa by analysis of microbiome composition with bias-correction (ANCOM-BC) (Supplementary Figure 2D,E). Taken together, this suggests that the OMM-12 community is not strongly altered by Mtb infection.

**Figure 2.**
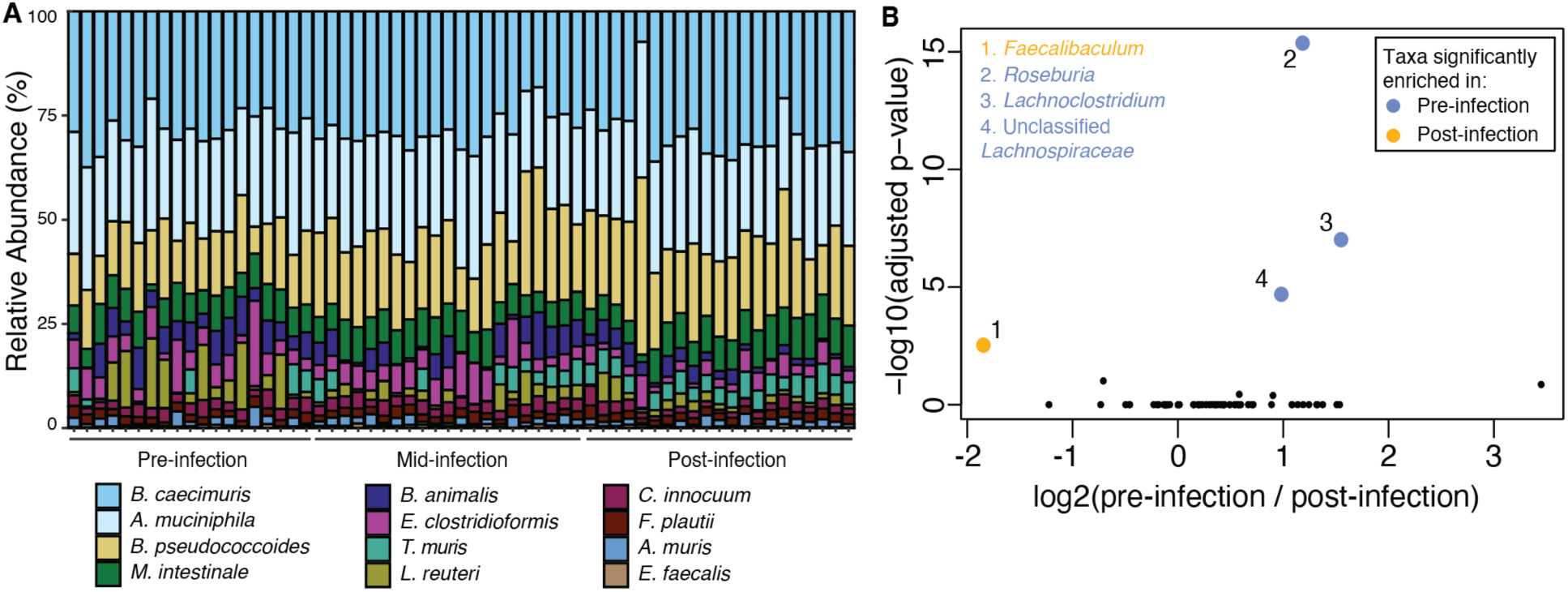
Mtb infection alters the microbiome composition of conventional MPF mice but not that of OMM-12 mice. (A) Taxonomic bar plot of the relative abundance of each OMM-12 taxa at days 0, 14, and 28 dpi (pre, mid, and post-infection respectively). Relative abundance is plotted at the genus level. (B) The log2(fold change) of each genera in MPF pre and post Mtb infection is plotted versus the log(adj p-value) as calculated by analysis of composition of microbiomes with bias correction (ANCOM-BC) (yellow and blue dots represent genera with an absolute log2(fold change) greater than 0.58 and an adjusted p-value less than 0.05).

In contrast, genus level analysis of the gut microbiome of MPF mice using ANCOM-BC revealed significant changes post-infection. *Faecalibaculum* species are enriched post-infection, whereas *Roseburia*, *Lachnochlostridium*, and Unclassified *Lachnospiraceae* species are depleted (Figure 2B, Supplementary Figure 3A-E). Additionally, subtle shifts in the alpha and beta diversities of these mice were observed pre- and post-infection (Supplementary Figure 3F-H).

### Gut colonization of *Helicobacter hepaticus* does not affect host susceptibility to Mtb

Previous studies in antibiotic treated immunocompromised mice correlated *H. hepaticus* colonization with a 0.75-fold log increase in Mtb lung burden and disease in mice with conventional microbiomes^15^. We sought to test this in wild type mice with a controlled microbiome background. Age matched OMM-12 and MPF mice were orally gavaged with the ATCC 51449 strain of *H. hepaticus* and colonization was confirmed via PCR on fecal gDNA. One week after *H. hepaticus* colonization, mice were intranasally infected with the Mtb H37Rv strain. At 28 days post Mtb infection, we measured lung bacterial burden and profiled immune changes (Figure 3A). In mice colonized with OMM12, the addition of *H. hepaticus* had no effect on Mtb burden in the lungs. In MPF mice, we saw that introduction of *H. hepaticus* caused a modest decrease in bacterial burden in the lungs of less than a 0.5 log fold, in contrast to what was previously reported in which *H. hepaticus* slightly enhanced susceptibility to infection. Overall, we found that *H. hepaticus* has only modest impacts on the outcome of Mtb infection in the setting of a controlled microbiome.

**Figure 3.**
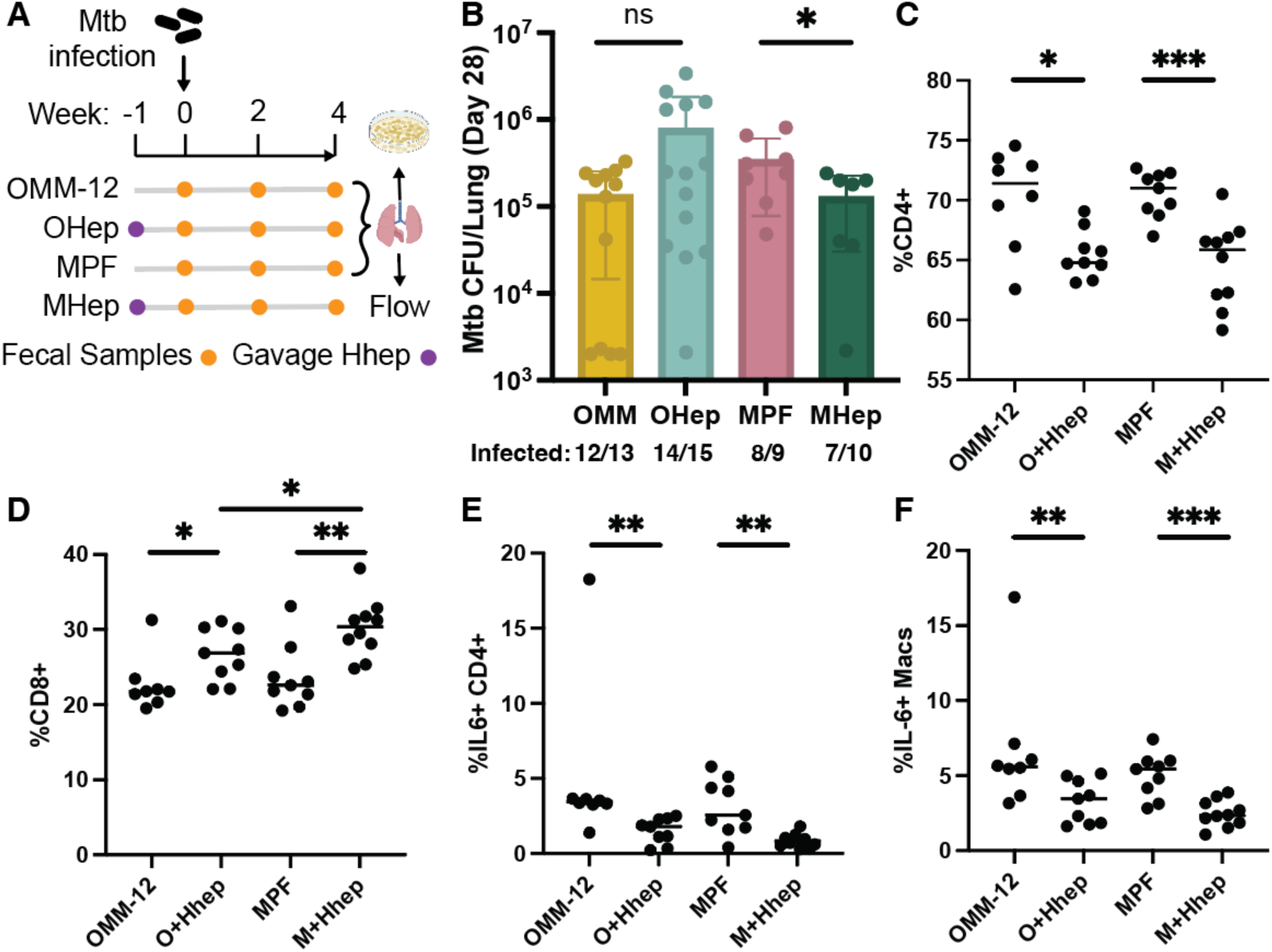
Addition of *H. hepaticus* to OMM-12 and MPF mice shows little difference in the susceptibility to Mtb, but shows a change in the immune microenvironment. (A) Experimental design of intranasal Mtb infections of *H. hepaticus* colonized mice and their control counterparts. *H. hepaticus* ATCC 51449 was orally gavaged into the mice at least 7 days prior to Mtb infection. Colonization was confirmed via PCR on fecal genomic DNA. Fecal samples were collected at 0, 14, and 28 dpi. Lungs were harvested at 28 dpi and used for plating for CFU (B) and flow cytometry (C-F). Immune cell populations and cytokines were measured after stimulation with ESAT6 (C) or Ag85B (D-F) peptide pools. Results in B-F are pooled from at least 2 independent experiments. *p < 0.05, **p < 0.01, ***p < 0.001, NS (Mann-Whitney U test).

### *H. hepaticus* colonization influences the lung immune microenvironment

We sought to evaluate the lung immune response to Mtb in the presence and absence of *H. hepaticus*, which has previously not been reported in detail. We performed a broad flow cytometry panel looking at innate and adaptive cell markers as well as cytokine markers indicative of *H. hepaticus* colonization. Antigen specific T cell responses were evaluated after stimulation of cells isolated from lungs with peptide pools of the Mtb T cell antigens ESAT6 and Antigen 85B (Ag85B). *H. hepaticus* colonization was associated with a decrease in CD4+ T cell populations (Figure 3C) and an increase in CD8+ T cell populations (Figure 3D). Cytokines relevant to Mtb disease burden (IL-10, IL-6, and TNFa) have been previously shown to be altered in the intestines of mice colonized with *H. hepaticus*^15,32^. We observed a decrease in IL-6 production from CD4+ T cells (Figure 3E) and macrophages (Figure 3F). We saw no differences in the production of IL-10 and TNF-α (Supplementary Figure 4). These results are the first demonstration that *H. hepaticus* colonization can alter the lung immune response to Mtb infection.

## Discussion

Humans have co-evolved with *M. tuberculosis* for millennia both before and after the age of widespread antibiotics. Tuberculosis disease progression is highly heterogeneous in the human population. No single factor can explain how and why tuberculosis severity varies between individuals. We sought to explore whether the gut microbiome can impact TB susceptibility using a new TB gnotobiotic mouse model. To our knowledge, germ-free mice have not previously been infected with Mtb. Current mouse models for analyzing the gut microbiome and pathogen interactions rely heavily on antibiotic treatment^7,15,33–36^. Antibiotics broadly disrupt the gut microbiome, providing a quick and simple way to study how general alterations of the gut microbiome can impact a particular disease state^37,38^. It also provides a feasible way to study gut microbiome alterations in high containment facilities such as the BSL3. Studies using antibiotic treatment have shown how particular taxa such as *Akkermansia muciniphila*^12^ and *Bacteroides fragilis*^13^ are beneficial in providing modest protection against Mtb via immune modulation in conventional mice. These taxa are also depleted in humans with active pulmonary TB^12,13^, further supporting their potential role in protection against tuberculosis. However, if the gut microbiome played a major role in susceptibility or protection to Mtb, we would expect to see increased susceptibility in germ-free mice. Surprisingly, we found that microbial diversity does not influence host control in an intranasal model of Mtb. Our OMM-12 community contains *Akkermansia* and *Bacteroides* species, yet we saw no significant difference in the susceptibility to Mtb in these mice compared to conventional or germ-free mice. Antibiotics may have systemic effects that impacted prior studies^39,40^. The addition of strains in a non-gnotobiotic setting might also have secondary impacts on other strains within the complex community. In antibiotic treatment models, it is not straightforward to attribute phenotypes solely to microbiome depletion or even addition of specific strains.

The effect of TB disease on gut microbiome composition has primarily been characterized in human studies comparing individuals with pulmonary TB to healthy controls^4–6^. Observed human microbiome composition changes correlated with disease are challenging to interpret since they could be a result of the disease, disease treatment, or the result of preexisting differences between individuals. In conventionally raised mice, we saw enrichment of *Faecalibaculum* and depletion of *Roseburia*, *Lachnoclostridium*, and unclassified *Lachnospiraceae* post TB infection. *Faecalibaculum* strains have previously been implicated in promoting depression^41^ and suppressing colorectal cancer^42^. The family *Lachnospiraceae*, for which *Roseburia* and *Lachnoslostridum* belong to, have been associated with both beneficial roles as SCFA producers^43^ and detrimental roles in multiple sclerosis^44^ and colorectal cancer^45^. Other studies looking at changes to the murine microbiome post Mtb challenge also demonstrated depletion of *Lachnospiraceae* upon aerosol infection of BALB/c mice^46^. Our findings add to the prior evidence that Mtb infection can alter the gut microbiome composition.

We also tested the effect of *H. hepaticus* colonization in our gnotobiotic model. *H. hepaticus* asymptomatically colonizes the gut of laboratory mice and causes chronic hepatitis and inflammatory large bowel disease in immunodeficient mice^47–49^. It also serves as a model to understand how microbiome-host interactions yield inflammatory and tolerogenic responses^50^. *H. hepaticus* induces IL-10 Treg^51^ and resident macrophage^32^ responses in the intestine, along with the production of IL-6 and TNF-α. Immune responses to *H. hepaticus* colonization have been characterized in the context of large intestine and related cancers, but the immune response at distal and barrier sites were unexplored prior to this study. Prior work established *H. hepaticus* colonization slightly increased susceptibility to Mtb and worsened immune response to Mtb in immunocompromised conventional B10.D2 mice^15^. However, we did not observe strong changes in Mtb lung burden driven by *H. hepaticus* colonization in either our defined community (OMM-12) or their conventional counterparts using C57BL/6 mice. Instead, we saw modest changes to the lung immune environment in mice colonized with *H. hepaticus*.

Germ-free and gnotobiotic mice allow for rigorous testing of gut microbiome effects on health and disease outcomes^52^, as gnotobiotic techniques allow for the complete control of gut microbial composition. Germ-free mice are more susceptible to some infectious diseases due to their poorly developed immune systems^53^. This phenotypic difference in disease burden has been shown in the context of *Listeria monocytogenes*^54^, murine cytomegalovirus and *Klebsiella pneumoniae* co-infections^55^. However, this phenotype seems to be pathogen- and organ-dependent. In the context of respiratory pathogens, germ-free mice show increased burden in the intestine during *K. pneumoniae* infection, but not in the lungs^56^. Our observation that germ-free mice are not more susceptible may be due to Mtb’s resistance to innate immunity^57^ and antimicrobial effectors^58–60^. Consistent with our findings, pet shop mice and conventional mice are similarly impacted by Mtb infection, despite very different gut microbiomes^61^.

Due to the technical complexity of this experimental setup, our study does have limitations. To keep the germ-free and gnotobiotic mice as sterile as possible in a BSL3 setting, we were limited to infecting mice intranasally. Intranasal administration of Mtb does not recapitulate the natural route of aerosol transmission of Mtb and leads to a broad range of CFU. Additionally, we had our OMM-12 and MPF mice colonized with *H. hepaticus* for just 1 week prior to Mtb challenge. Longer colonization might lead to phenotypic differences in lung burden as it would give *H. hepaticus* more time to prime the immune response. However, even with 1 week of *H. hepaticus* colonization we were able to see detectable differences in the lung immune response to Mtb (Figure 3c-f).

In conclusion, we find that gut microbiome composition does not impact Mtb lung burden, regardless of complexity (germ-free versus OMM-12 versus conventional). Even when the lung immune environment is perturbed by *H. hepaticus* colonization, Mtb colonization is not affected. Instead, Mtb infection causes changes in the gut microbiome. It remains to be seen if this same directionality of interaction is at play in humans, or whether gut dysbiosis might impact severity of disease or response to treatment. Our findings highlight the challenging nature of TB and its ability to circumvent the mammalian immune system. Identifying new therapeutic avenues to prevent and cure TB remains a pressing global issue.

## Funding

This study was supported by the National Institutes of Health (R25AI147375, R35GM147512, R21AI176295, 1R01AI153197-01, P01AI063302). CA was supported by Gilead Sciences, NSF GRFP, and the Hellman Foundation.

## Acknowledgements

We thank members of the Wolf and Stanley labs for their feedback and support. We would like to thank the Cox and Vance labs for the help and advice received, as well as the Barton lab for their help with the OMM-12 community. We additionally thank Jessica Hoisington-Lopez and MariaLynn Crosby at Washington University in Saint Louis for sequencing support.

## Supplement

**Supplementary Table 1.** Oligo Mouse Microbiota 12 Strains.

| Taxonomic Classification - Phylum | Strain Name | DSMZ Strain ID |
| --- | --- | --- |
| Actinomycetota (Actinobacteria) | *Bifidobacterium animalis* | YL2 |
| Bacillota (Firmicutes) | *Acutalibacter muris* | KB18 |
|  | *Blautia pseudococcoides* | YL58 |
|  | *Clostridium innocuum* | I46 |
|  | *Enterocloster clostridioformis* | YL32 |
|  | *Enterococcus faecalis* | KB1 |
|  | *Flavonifractor plautii* | YL31 |
|  | *Limosilactobacillus reuteri* | I49 |
| Pseudomonadota (Proteobacteria) | *Turicimonas muris* | YL45 |
| Bacteroidota | *Bacteroides caecimuris* | I48 |
|  | *Muribaculum intestinale* | YL27 |
| Verrucomicrobiota | *Akkermansia muciniphila* | YL44 |

**Supplementary Table 2.**
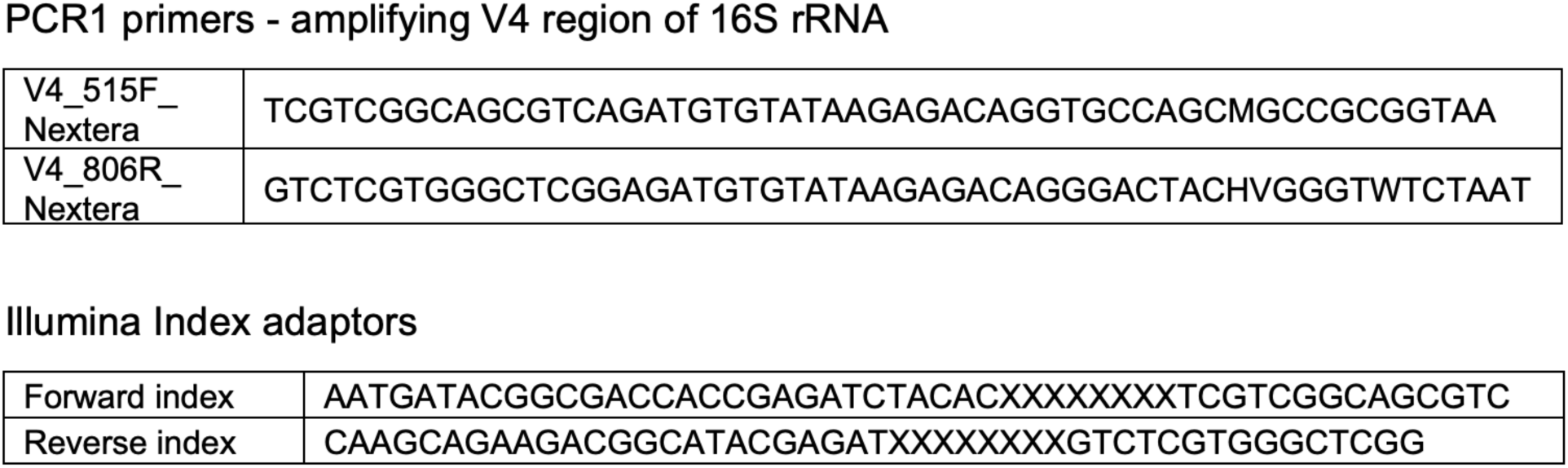
Primer and adaptor sequences.

**Supplementary Figure 1.**
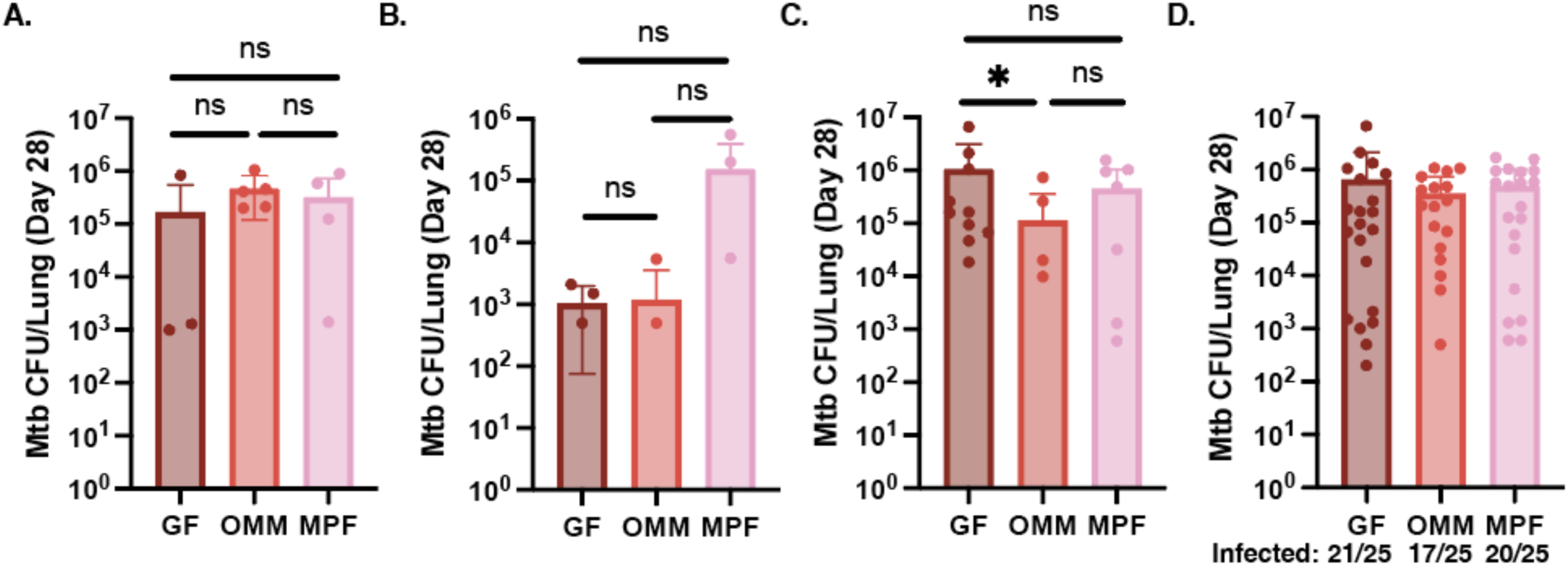
Mtb susceptibility is independent of gut microbiome diversity. Lungs were harvested at 28 dpi and used for plating for colony forming units (CFU). (A-C) Mtb burden in the lung determined by CFU counts at 3 weeks post harvest across 3 independent experiments (NS, Mann-Whitney U test). (D) Combined Mtb burden across all experiments. Number of infected mice out of total attempted infections is shown below.

**Supplementary Figure 2.**
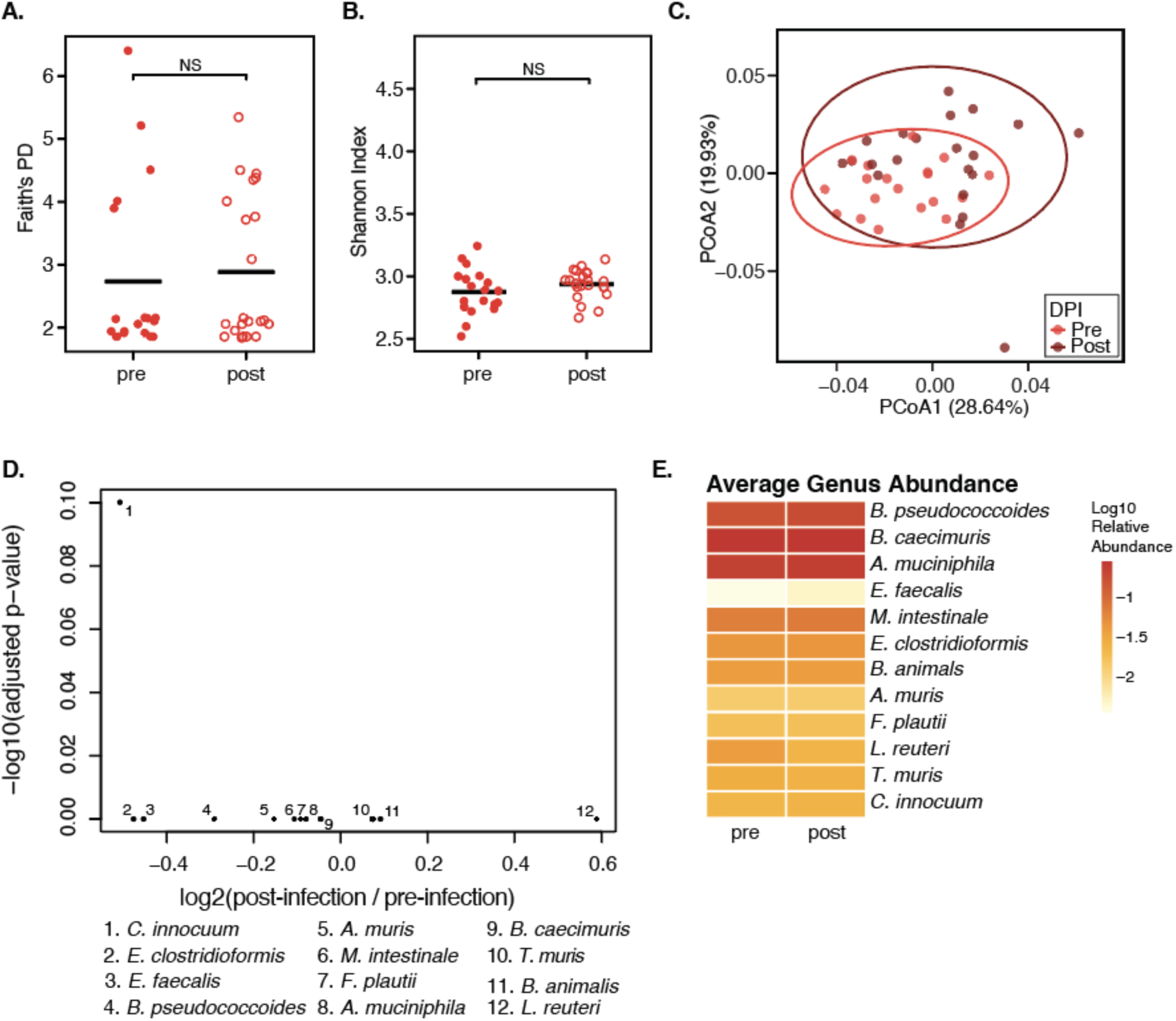
Mtb infection does not affect the OMM-12 community in the intestine. Alpha diversity of OMM-12 mice pre and post Mtb infection measured by (A) Faith’s PD and (B) Shannon Index (Wilcoxon rank sum test, NS). (C) Principal coordinate analysis (PCoA) calculated by Weighted UniFrac distance (p < 0.01, PERMANOVA; p = 0.412, PERMDISP). (D) ANCOM-BC on OMM-12 post-infection over pre-infection (log2-fold change < 0.58, padj < 0.05, NS). (E) Average genus abundance (log10 relative abundance) of each member of the OMM-12 community.

**Supplementary Figure 3.**
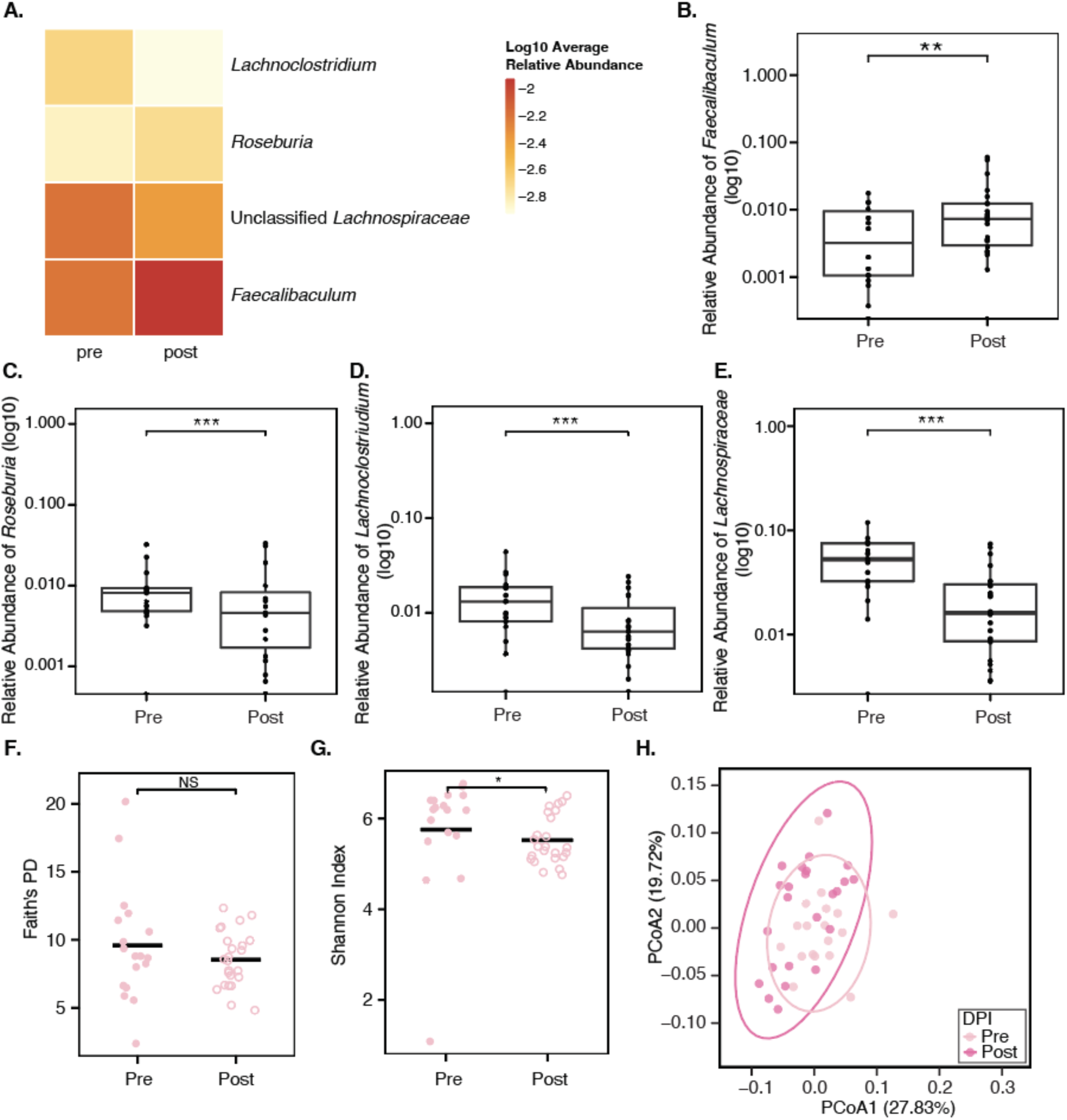
Mtb infection leads to gut microbiome alterations in MPF mice. (A) Average genus abundance (log10 relative abundance) of each genus identified as differentially abundance by ANCOM-BC. Box plots of the relative abundances (log10) of (B) *Faecalibaculum*, (C) *Roseburia*, (D) *Lachnoclostridium*, and (E) Unclassified *Lachnospiraceae* (padj** < 0.01, padj*** < 0.001). Alpha diversity of MPF mice pre and post Mtb infection measured by (F) Faith’s PD and (G) Shannon Index (WIlcoxon rank sum test, NS, p* < 0.05). (H) PCoA calculated by Weighted UniFrac distance (p < 0.01, PERMANOVA; p = 0.945, PERMDISP).

**Supplementary Figure 4.**
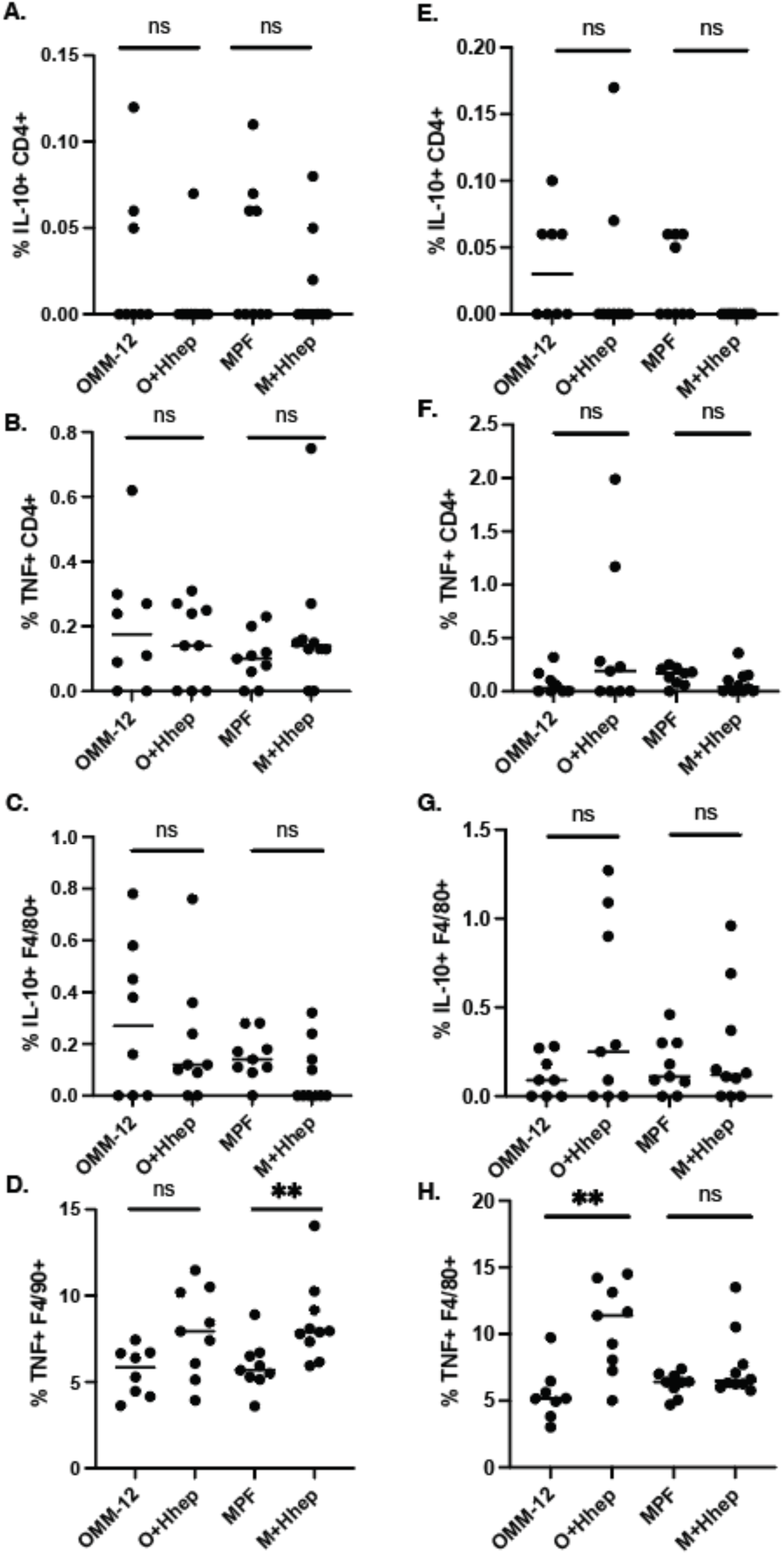
H. hepaticus-associated cytokines are not impacted by Mtb infection. *H. hepaticus* cytokines were measured after stimulation with ESAT6 (A-D) or Ag85B (E-H) peptide pools. Results are pooled from at least 2 independent experiments. **p < 0.01, NS (Mann-Whitney U test).

## Notes

### Competing Interest Statement

The authors have declared no competing interest.

